# Loss of G9a does not phenocopy the requirement for Prdm12 in the development of the nociceptive neuron lineage

**DOI:** 10.1101/2023.09.15.557867

**Authors:** Panagiotis Tsimpos, Simon Desiderio, Pauline Cabochette, Sadia Kricha, Eric J. Bellefroid

**Affiliations:** ULB Neuroscience Institute (UNI), Université Libre de Bruxelles (ULB), Gosselies, Belgium

**Keywords:** dorsal root ganglia, neurogenesis, somatosensory neurons, nociceptors, G9a, Prdm12

## Abstract

Prdm12 is an epigenetic regulator expressed in developing and mature nociceptive neurons, playing a key role in their specification during neurogenesis and modulating pain sensation at adulthood. *In vitro* studies suggested that Prdm12 recruits the methyltransferase G9a through its zinc finger domains to regulate target gene expression, but how Prdm12 interacts with G9a and whether G9a plays a role in Prdm12’s functional properties in sensory ganglia remain unknown. Here we report that the SET domain of G9a is necessary and sufficient for the interaction with Prdm12. We show that Prdm12 is co-expressed with G9a in dorsal root ganglia during early murine development. To address the role of G9a in somatosensory neurogenesis and test the hypothesis that it may function as a mediator of Prdm12’s function during somatosensory neurogenesis, we conditionally inactivated it in neural crest using a Wnt1-Cre transgenic mouse line. We found that G9a ablation in neural crest does not lead to dorsal root ganglia hypoplasia due to the loss of somatic nociceptive neurons nor to the ectopic expression of the visceral determinant Phox2b as observed upon *Prdm12* ablation. Together, our results confirm Prdm12’s ability to interact with G9a and reveal that this interaction is however not instrumental for its developmental function during nociceptive neuron development.

## INTRODUCTION

The bodily ability of vertebrates to discriminate and respond to a wide array of salient stimuli resides in the great diversity of neuron subtypes whose cell bodies are found in dorsal root ganglia (DRG) and build a discriminative sensory relay between the periphery and the central nervous system. This neuronal diversity arises during development when specific transcriptional programs are initiated that bias the fate of neural crest-derived somatosensory progenitors into one of the three cardinal somatosensory lineages. These three main subtypes of somatosensory neurons can be discriminated early in development based on the selective expression of tyrosine kinase neurotrophic receptors [1–4]. Tyrosine kinase receptor A (TrkA)-is expressed in developing lightly or unmyelinated nociceptive neurons of small or medium diameter which mostly respond to noxious stimuli but are also involved in temperature or itch sensing [5] and in unmyelinated low-threshold mechanoreceptive (LTMR) neurons involved in pleasurable touch [6–8]. Ret, TrkB and TrkC are expressed in more myelinated LTMR neurons that convey innocuous touch sensation and proprioception. These early developmental selective expressions eventually evolve over time as sensory neurons mature and diversify into more specialized subtypes, with the wider diversification seemingly arising in the TrkA lineage [3,9].

Over the last decades, a comprehensive understanding of the main transcriptional regulators guiding the early development and diversification of somatosensory neurons has been acquired. Notably, molecular players required for the emergence and diversification of the TrkA lineage have been identified [3,4]. Among them, the transcriptional regulator Prdm12 stands at the root of the specification of this lineage. Indeed, Prdm12 function is critical for the emergence of the entire pool of neurons arising from immature TrkA-expressing sensory neuron precursors as well as for TrkA expression itself [10–13]. Accordingly, in human, patients harbouring homozygous mutations of *PRDM12* suffer from congenital insensitivity to pain (CIP), a rare developmental disorder associated with depletion of somatosensory fibers allowing the detection of noxious stimuli [14,15]. Moreover, as *Prdm12* remains expressed in mature nociceptive neurons and its induced conditional ablation at adulthood alters some pain-related behaviours, it has been hypothesized as a potential new therapeutic target to treat pain related diseases [10,16,17]. However, to approach the therapeutic potential of Prdm12, a deeper understanding of its mode of action needs to be first established.

Prdm12 is a member of the PRDM family of epigenetic zinc-finger transcriptional regulators playing roles in many developmental processes and diseases [18–21]. While some PRDM family members possess intrinsic histone methyltransferase activity through their SET related PR domain, it appears not to be the case for Prdm12 which has been shown to be able to form a complex when overexpressed in H2K29T3 cells with G9a, a histone methyltransferase (HMT) that dimethylates histone H3 at lysine 9 (H3K9me2), a hallmark of epigenetic repression [22,23]. While some evidence have been obtained using P*rdm12* overexpression experiments in *Xenopus* suggesting that this interaction with G9a may be functionally relevant for Prdm12’s activity [15,23], no *in vivo* evidence supporting this hypothesis is available.

In this study, using co-immunoprecipitation assays in HEK293T cells, we show that Prdm12 interacts with the SET domain of G9a. To evaluate G9a function in sensory neurogenesis and possible importance for Prdm12 functions in developing DRG, we generated *G9a* conditional knockout murine embryos (*G9a* cKO) in which *G9a* is selectively invalidated in the neural crest lineage. We report here that the loss of G9a does not phenocopy the requirement of Prdm12 for the initiation of the nociceptive lineage. These data suggest that G9a is not instrumental for Prdm12 function during somatosensory neurogenesis.

## RESULTS

### Prdm12 interacts via its zinc fingers with the SET domain of G9a

Previous studies have highlighted the ability of Prdm12 to interact with G9a. To define which region of G9a is responsible for the interaction with Prdm12, Myc-G9a deletion mutants were generated that eliminate some of the conserved domains of the protein (MYC-G9a ΔSET, ΜYC-G9a ΔΑΝΚ, MYC-G9a SET). Constructs encoding these mutants were co-transfected in HEK29T3 cells together with a construct encoding a Flag tagged version of mPrdm12. In co-immunoprecipitation (Co-IP) assays, we found that while the MYC-G9a ΔANK and MYC-G9a SET proteins bind to Flag-Prdm12, MYC-G9a ΔSET was unable to do so **(Figure 1A, B)**. As reported previously [22], a mutant version of mPrdm12 lacking the ZF domain was unable to interact with G9a. Thus, Prdm12 interacts via its zinc finger domain with the SET domain of G9a.

**Figure 1.**
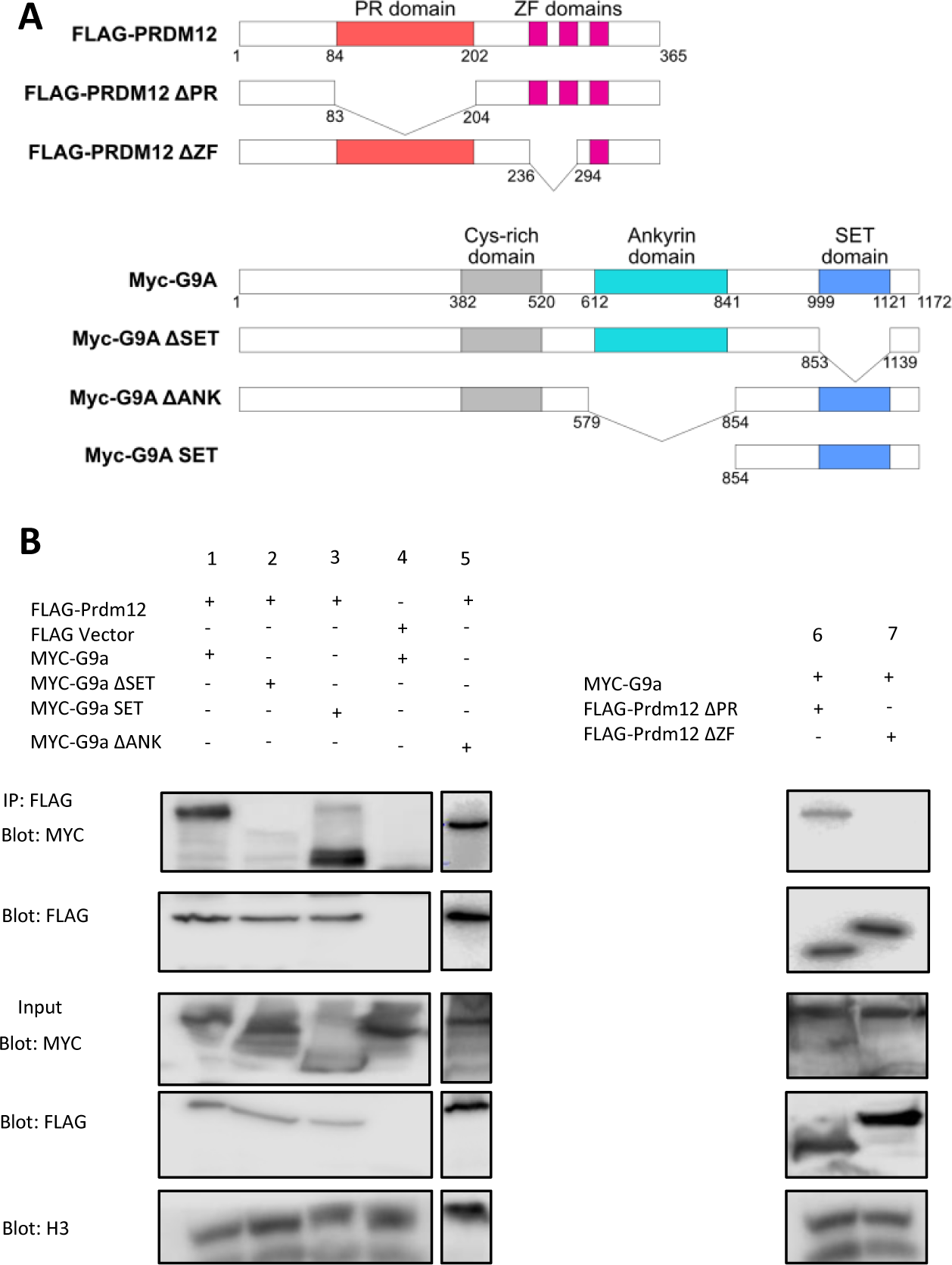
Prdm12 interacts via its zinc fingers with the SET domain of G9a. (A) Schematic diagram of WT and deletion mutants of Flag-PRDM12 and Myc-G9A, using an empty FLAG vector as a control. **(B)** HEK293T cells were transfected with the indicated plasmids. Lysates were immunoprecipitated with anti-Flag antibodies. Immunoprecipitates and 5% of the input were then subjected to western blot analysis with anti-Flag or anti-Myc antibodies.

### G9a and Prdm12 are co-expressed during somatosensory neurogenesis

As a first step to test the hypothesis that Prdm12 functions through G9a to regulate development of the TrkA lineage during somatosensory neurogenesis, we first compared by immunofluorescence the expression of G9a and Prdm12 in DRG of murine embryos from E10.5 to E12.5. **Figure 2A-B** shows that G9a is like Prdm12 broadly expressed in DRG at these stages. While all Prdm12^+^ cells colocalized with G9a, some G9a^+^/Prdm12^-^ cells were also clearly visible (**Figure 2C**). Thus, G9a is broadly expressed during somatosensory neurogenesis and is co-expressed with Prdm12 in the developing nociceptive neuron lineage, which suggest that they may functionally interact to control its development.

**Figure 2.**
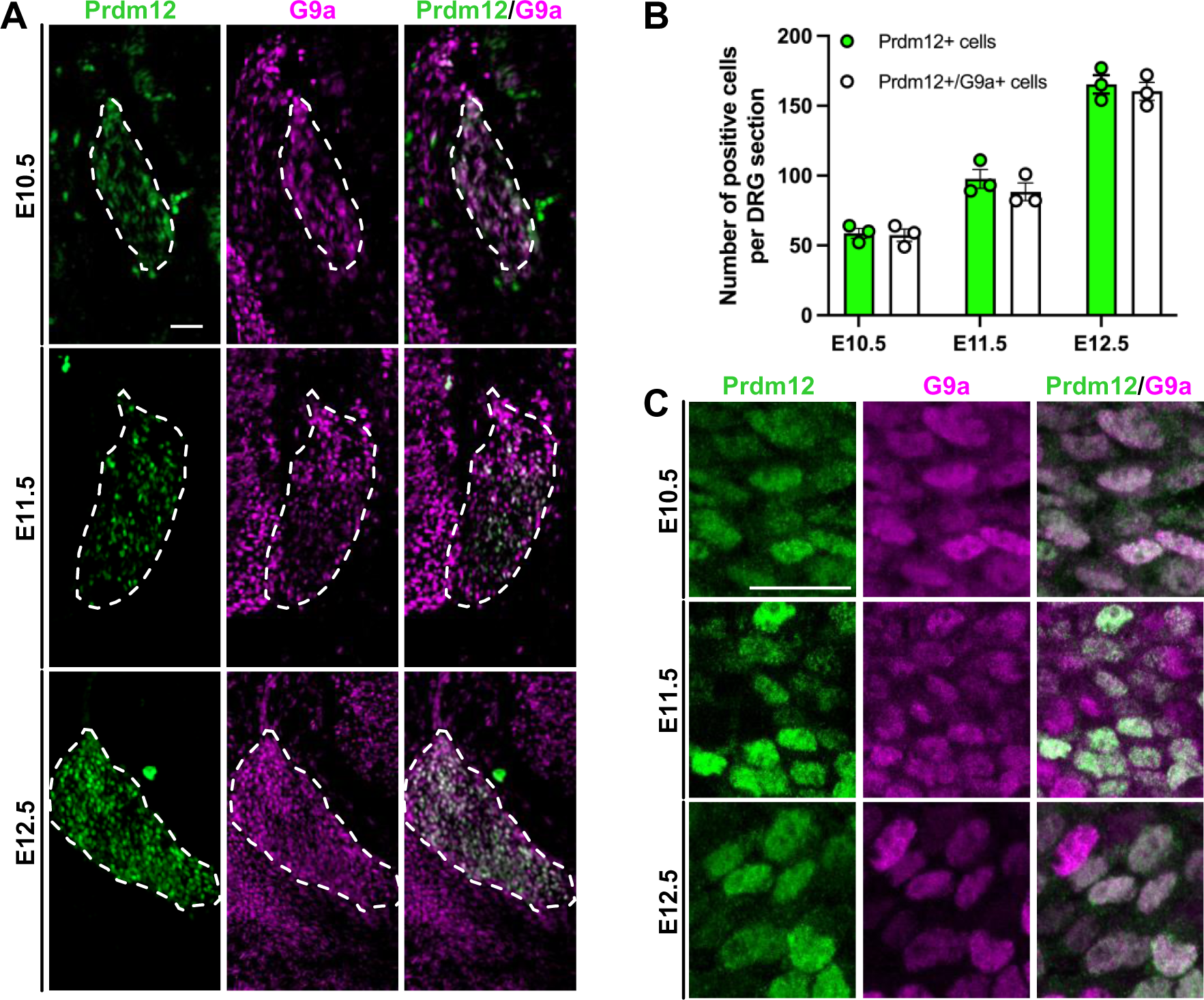
G9 is coexpressed with Prdm12 in developing dorsal root ganglia. (A) Immunostainings for Prdm12 and G9a on coronal sections through DRG of wild-type mouse embryos at indicated stages. DRG are delineated by white dashed lines. Scale bar, 50 µm. **(B)** Quantification of the mean number of Prdm12^+^ cells or Prdm12^+^/G9a^+^ cells numbered in DRG coronal sections of wild-type embryos at indicated stages. Histograms are represented as mean ± SEM. Each dot represents the mean value obtained for an individual biological replicate. **(C)** High magnification pictures of representative immunostainings as described in panel A. Scale bar, 50 µm.

### Neural crest specific depletion of G9a does not recapitulate the requirement of Prdm12 for genesis of the TrkA-lineage in embryonic DRG

If G9a is a key mediator of Prdm12’s function, then its loss should recapitulate the phenotype observed in Prdm12 knock-out mice. To test this hypothesis, we generated a *G9a* conditional knockout (cKO) mouse line by crossing *G9a* homozygous floxed mice (exons 26-28) with mice carrying a transgene expressing the Cre-recombinase under the control of the neural crest specific *Wnt1* promoter [24,25]. The *G9a* cKO mice generated from this strategy survived until late development (E18.5) but never thrived in the mouse litters. Immunofluorescence analysis of G9a expression in E12.5 embryos confirmed its selective depletion in DRG (**Figure S1A-B**). RT-qPCR analysis performed on control and *G9a* cKO E14.5 isolated DRG further confirmed the efficiency of the deletion (**Figure S1C**). H3K9me2, that is catalyzed by G9a was as expected dramatically reduced in *G9a* cKO compared to controls. In contrast, H3K9me2 was not lost in *Prdm12* KO embryos (**Figure S1D**), suggesting that Prdm12 does not rely on H3K9me2 marks to mediate its function or that the loss of *Prdm12* alone, which is only one of the putative partners of HMT proteins, is not sufficient to observe a reduction of staining. Together, these data indicated that *G9a* is efficiently deleted in our *G9a* cKO.

In developing DRG, *Prdm12* is selectively expressed in the TrkA lineage which accounts for the vast majority of DRG somatosensory neurons. Following *Prdm12* knockout this whole lineage fails to develop, concomitantly resulting in a dramatic hypoplasia of DRG [10–13]. To assess the role of G9a in DRG neurogenesis, we first performed immunofluorescence staining using the pan-sensory neuronal marker Islet1 on coronal sections through DRG of E11.5 to E14.5 control and *G9a* cKO embryos. No difference in the number of Islet1^+^ neurons was found between *G9a* cKO and control embryos, at any of the stages examined (**Figure S1E**). No difference was also observed between *G9a* cKO and control embryos at E11.5, E12.5 and E14.5 in the number of nociceptive neurons as visualized by TrkA or Prdm12 immunostaining (**Figure 3A-C**). This contrasts with the severe neuronal loss and DRG hypoplasia observed in *Prdm12* KO embryos from E13.5 onward following the agenesis of the TrkA neuron lineage [12].

**Figure 3.**
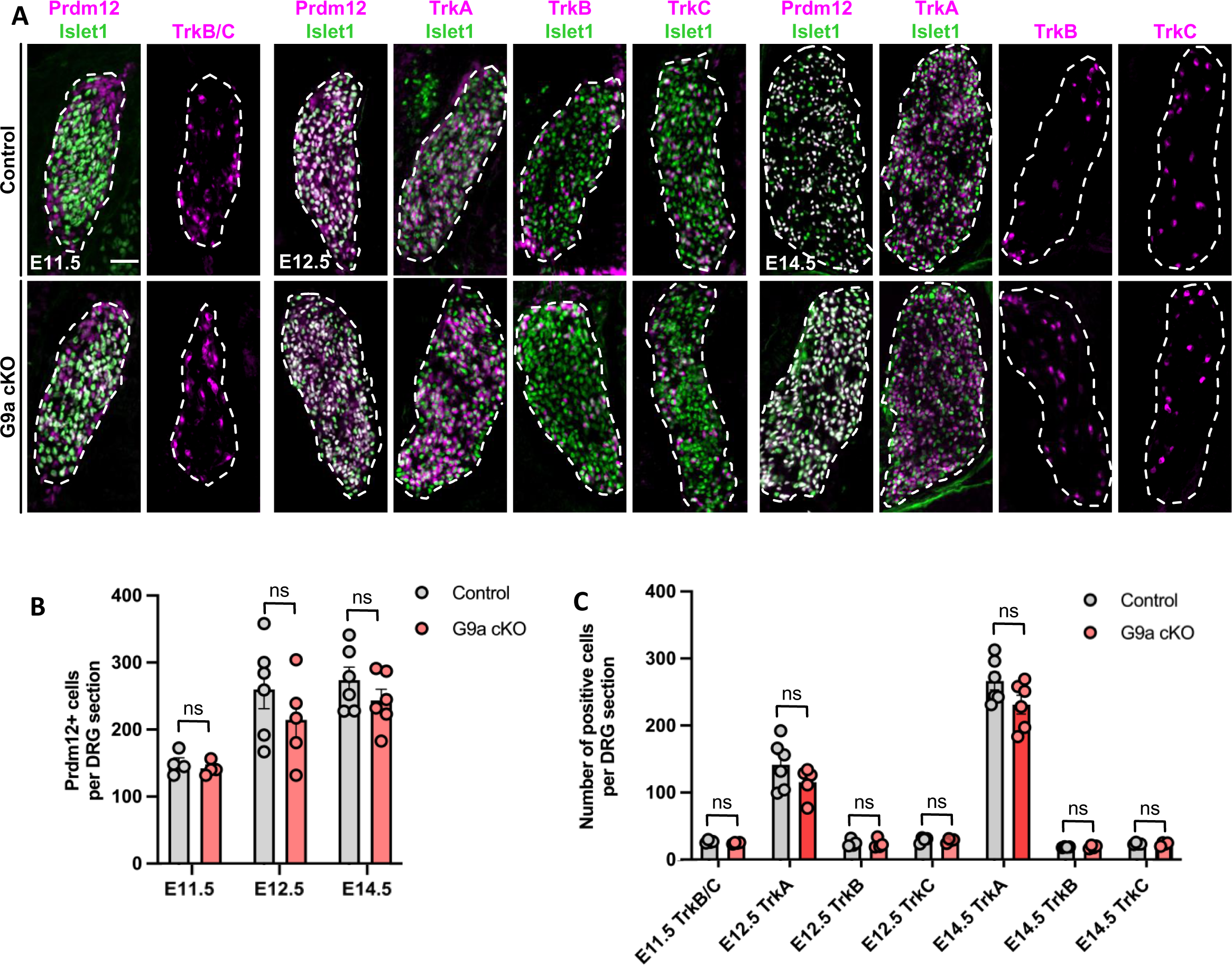
Loss of G9a is dispensable for early sensory neuron development in dorsal root ganglia. (A) Double immunostainings with indicated markers on coronal sections through DRG of control or *G9a* cKO embryos at E11.5, E12.5 and E14.5. Scale bars, 50 µm. DRG are delineated by white dashed lines. **(B-C)** Quantification of the mean number of Prdm12^+^ cells (A) or cells positive for TrkA, TrkB or TrkC (B) in DRG coronal sections of control or *G9a* cKO embryos at indicated stages. Histograms are represented as mean ± SEM. Each dot represents the mean value obtained for an individual biological replicate. Mann-Whitney test. P-value, ns > 0.999.

In embryonic DRG of *Prdm12* KO embryo, increased apoptosis and reduced cell proliferation have been reported to contribute to the agenesis of the TrkA neurons lineage [11–13]. Therefore, and given the importance of G9a in the control of cell proliferation and survival [26], we also investigated the consequences of the loss of *G9a* on cell death using immunostaining with anti-activated caspase 3 antibodies and on cell proliferation using phospho-Histone H3 antibodies. While no cell proliferation defect was detected in *G9a* cKO, a transient increase of apoptosis was found at E11.5, which was not detectable anymore at E14.5 (**Figure S2**). Thus, in *G9a* cKO embryos, a transient increase of apoptosis appears to occur that however does not result in the loss of the nociceptive lineage, as observed in *Prdm12* KO [13].

Phox2b is a master regulator of visceral fates in the peripheral nervous system [4,27,28]. As we recently discovered that Prdm12 also promotes nociceptor fate by repressing Phox2 genes and thus preventing precursors from engaging into an alternate visceral neuronal differentiation program (accepted in iScience), we also analyzed Phox2b expression in DRG of *G9a* cKO embryos. While Phox2b positive cells were indeed observed in E11.5 *Prdm12* KO DRG, none could be detected in *G9a* cKO (**Figure S3**).

As G9a appears more broadly expressed than Prdm12 in embryonic DRG, we next also examined the consequence of the loss of *G9a* on the non-nociceptive neuron lineages, following the expression of the neurotrophic receptors TrkB and TrkC, labelling mechano/proprioceptive neurons. No difference was again observed using these markers between *G9a* cKO and controls (**Figure 3A-C**). Together, these results indicate that G9a is not essential for somatosensory neurogenesis and that it does not act as a critical mediator for Prdm12 functions in the initiation of the nociceptive neuron lineage.

### G9a appears dispensable for the maturation of the three main subtypes of somatosensory neurons

Sensory neurogenesis in DRG begins around E9.5 and is complete by E14.5. It further overlaps and is followed by a phase of developmental maturation during which somatosensory neurons further refine into more specialized somatosensory subtypes [1,3,9]. To determine if G9a is involved in this sensory fate refinement phase, we examined the expression of sensory subtype markers at E18.5. At this stage, TrkB-expressing neurons label subtypes of myelinated LTMR neurons while TrkC is also found in LTMR subtypes and in proprioceptors [9]. The nociceptive component of the TrkA lineage also undergoes a functional refinement with the gradual emergence of peptidergic (PEP) nociceptors, which maintain TrkA and begin to express neuropeptides such as Calcitonin-Gene Related Peptide (CGRP), and non-peptidergic (NP) nociceptors which express the neurotrophic receptor Ret and initiate a phase of TrkA extinction [29]. Most neurons of the TrkA lineage also start to express the sodium channel Nav1.8 as a critical component driving their mature electrophysiological properties. We performed immunostainings using antibodies against all above cited markers as well as against Prdm12 and the pan-sensory neuron marker Islet1 to get clues of any maturational discrepancy of somatosensory subtypes in E18.5 control and *G9a* cKO DRG (**Figure 4A-B**). However, again, no significant difference was observed between *G9a* cKO and controls. Thus, G9a appears also dispensable for the maturation of the three main subtypes of somatosensory neurons.

**Figure 4.**
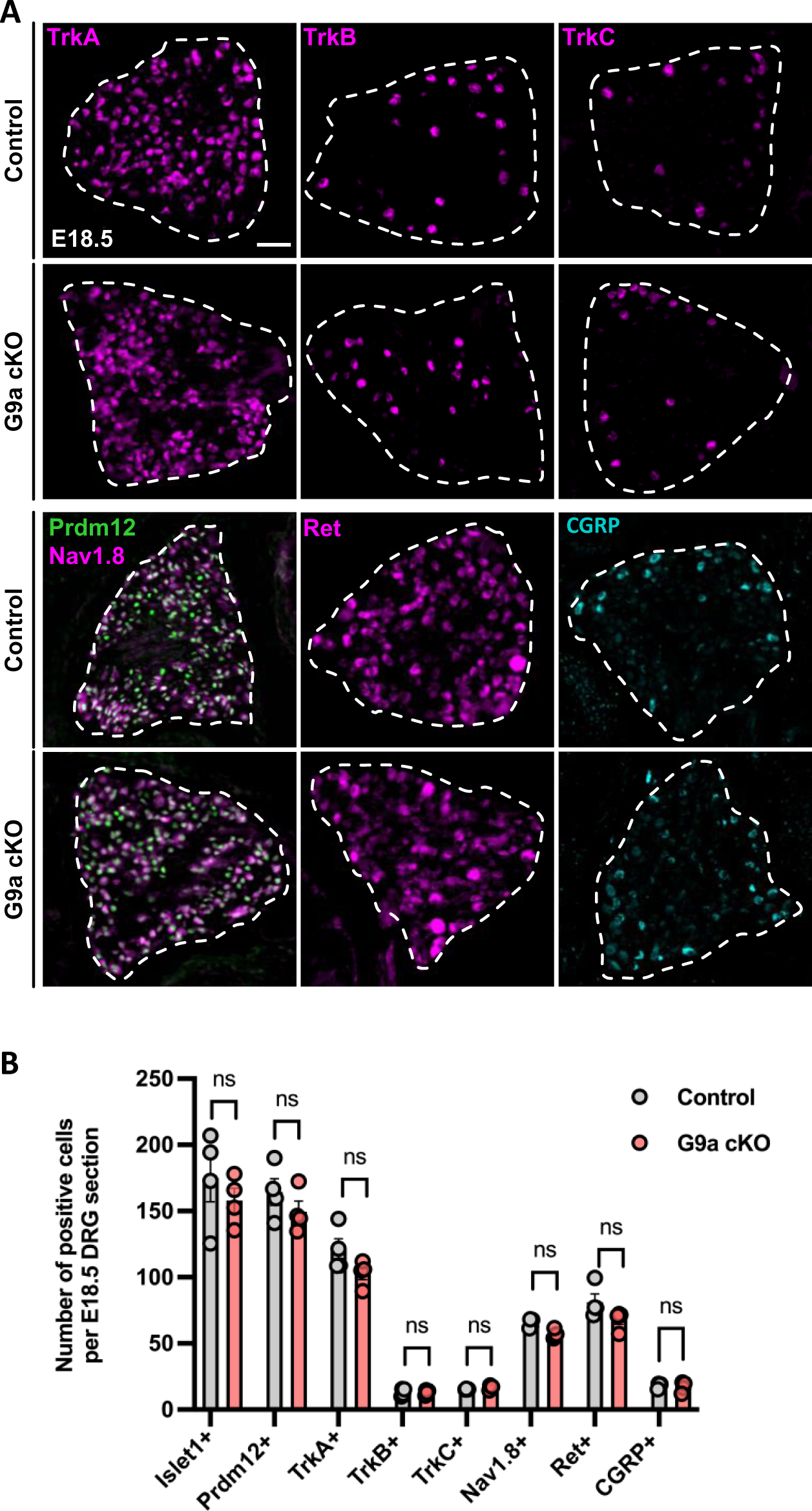
Loss of G9a is dispensable for the maturation of dorsal root ganglia sensory neurons. (A) Immunostainings with indicated markers of sensory neuron subtypes performed on coronal sections through DRG of control or *G9a* cKO embryos at E18.5. Scale bars, 50 µm. DRG are delineated by white dashed lines. **(B)** Quantification of the mean number of cells labelled with indicated markers on coronal sections through DRG of control or *G9a* cKO embryos at indicated stages. Histograms are represented as mean ± SEM. Each dot represents the mean value obtained for an individual biological replicate. Mann-Whitney test. P-value, ns > 0.999.

## DISCUSSION

PNS somatosensory neurons develop into specific subtypes thanks to the selective expression of a cascade of transcriptional regulators providing them with a discriminative transcriptional identity ultimately reflected by their subtype-specific functional diversity [3,5]. At the root of the TrkA-lineage, from which emerge most if not all small diameter somatosensory neurons (i.e, nociceptors and C-LTMR), stands the histone methyltransferase (HMT) related transcriptional regulator Prdm12. In the recent years, several studies have shown how much Prdm12 is instrumental in the emergence of the TrkA-lineage, which fails to develop in its absence [10–13,15]. Mechanistically, Prdm12 appears to act as a pseudo methyltransferase as it has been shown to interact with G9a when overexpress in HEK29T3 cells and to increase H3K9me2 level when overexpressed in *Xenopus* neuralized animal cap explants [15,22]. However, the *in vivo* relevance of G9a for Prdm12 functions in the emergence of the nociceptive lineage during somatosensory neurogenesis has never been investigated.

Here, using co-immunoprecipitation experiments, we confirm the ability of mPrdm12 to interact with G9a and that its first two ZF domains are required for this interaction. By contrast, Yildiz et al., 2019 found that the ZF domain of the zebrafish ortholog of Prdm12 is not required for this binding. The origin of this discrepancy is unclear as the sequence of the two first zinc finger domains of the mouse and zebrafish Prdm12 proteins is identical. We further demonstrate that the SET domain of G9a, that is responsible for its H3K9 methyltransferase activity and mediates homo-or heterodimerization with the related G9a-like protein (GLP) [25,26] is required and sufficient for the interaction with Prdm12. Like Prdm12, several other members of the Prdm family, including Prdm1, Prdm4, Prdm5, Prdm6 and Prdm16 have been shown to function as indirect epigenetic regulators and to be able to recruit G9a [30–33]. Whether the SET domain of G9a is also responsible for its ability to interact with these Prdm proteins remains to be determined.

We show that G9a and Prdm12 are co-expressed in embryonic DRG suggesting that G9a may play a role in Prdm12’s function in somatosensory neurogenesis. Although a transient increase of apoptosis was detected at E11.5, surprisingly our results of the analysis of the phenotype in DRG of G9a early cKO embryos revealed no deficiencies comparable to that of *Prdm12* KO embryos, suggesting the existence of an unknown catch-up mechanism. Specifically, in DRG of G9 cKO early embryos, no vanishing of the TrkA-lineage nor ectopic Phox2b expression was detected. At E18.5, no defect in the expression of late nociceptive markers was detected. As Prdm12 is unable to interact with GLP [22], and because the loss of G9a alone is sufficient to strongly reduce H3K9me2 level in DRG, it is unlikely that the lack of phenotype in DRG of G9a cKO embryos is due to compensation by GLP. We thus conclude that G9a is not required for Prdm12 function in the initiation of the nociceptive neuron lineage and is largely dispensable for trunk somatosensory neurogenesis. This contrast with its important role in cranial neural crest in bone formation [34,35].

G9a is also expressed in mature nociceptors and plays an important role in neuropathic pain [36]. Whether G9a functionally interacts with Prdm12 in this process remains a hypothesis to be investigated. How Prdm12 controls the expression of its targets during somatosensory neurogenesis remains thus today largely unknow. Prdm12 binds also to the histone methyltransferase EZH2 (Biogrid and **Figure S4**), a component of the polycomb repressive complex 2 but its loss in neural crest also does not interfere with sensory neuronal differentiation, suggesting it is also not instrumental for Prdm12 function [37]. Defining Prdm12’s interactome in an unbiased manner will thus be of paramount importance to understand its mechanism of action.

## DECLARATIONS

### Ethics approval

The animal experimental procedures were approved by the CEBEA (Comité d’éthique et du bien-être animal) of the IBMM-ULB and conformed to the European guidelines on the ethical care and use of animals.

### Consent for publication

Not applicable

### Availability of data and materials

All data used in the preparation of this manuscript will be provided upon request.

### Authors contributions

PT performed the molecular and biological experiments, with the help of SK, PC and SD. PT and SD analyzed the data and wrote the manuscript, with input from all other authors. EJB designed and supervised the study.

## Acknowledgements

We thank Dr. Yoichi Shinkai and Maite G. Fernández-Barrena for the *G9a* floxed mice and Dr. Alexandre Pattyn for the *Wnt1-Cre* mice. We thank Dr. Jean-François Brunet for providing the Phox2b antibody. We thank Marion Santangelo and Caroline Vanhulle for help with transfections and Louis Delhaye for assistance in the maintenance of mice at the mouse facility.

## MATERIAL AND METHODS

### Mouse ethics and crossing strategy

All mice were maintained on a C57BL/6J background. Mice were housed at room temperature with a 12h light/dark cycle in standard cages with litter, water and food *ad libitum*. Air circulation in the facility was filtered and temperature monitored at a steady 20°C. Cages were also provided with cottons and cardboard rolls for enrichment. The experimental protocols were approved by the CEBEA (Comité d’éthique et du bien-être animal) of the IBMM-ULB and are conform to the European guidelines on the ethical care and use of animals. The following mice strains were used: *Prdm12^LacZ/LacZ^* [13]*, Ehmt2/G9a^fl/fl^* (kindly gifted by prof. Maite García Fernández de Barrena, Universidad de Navarra)[25]*, Wnt1^Cre^*[38].

For the generation of the *Wnt1^Cre^; G9a^fl/fl^*mouse line (*G9a* cKO), *Wnt1^Cre^* males and females were crossed with *G9a^fl/+^* or *G9a^fl/fl^* females and males, respectively. *Wnt1^Cre^; G9a^fl/+^*mice were then crossed with *G9a^fl/fl^* mice to maintain the line and obtain control (*G9a^fl/+^, G9a ^fl/fl^*or *Wnt1^Cre^; G9a^fl/+^*) and *G9a* cKO embryos. For embryo harvesting, the day of vaginal plug was considered to be embryonic day (E) 0.5. A minimum of 8 sections per tissue and at least 4 embryos of the same genotype were analyzed in each experiment. Embryos were collected at E10.5, E11.5, E12.5, E14.5 and E18.5.

Polymerase Chain Reaction (PCR) was used for genotyping of the collected embryos as follows: For the *G9a* floxed and WT alleles, using primers forward 5’-CTGCACGCTGCCTAGATGGAGCATG-3’ and reverse 5’-CTGGGTGGAAAGTTGCCAGGCTTAG-3’, for the *Wnt1^Cre^* transgene, using primers forward 5′-CCACCTCTTCGGCAAGATCG-3′ and reverse 5′-GCTAGAAAGAATCTGGTGCTGACC-3′, for the *Prdm12^LacZ^* and WT alleles, using primers forward 5’-AGTTTGTACATTCCCTGGGAGTAAGACTCC-3’ and reverse 5’-AGCCAGGGGAAGAATGTGAGTTGC-3’.

### Plasmids and cloning

Flag-mPrdm12 WT and mutant and G9a (S) expression plasmids were kindly provided by Prof. Yoichi Shinkai, University of Kyoto, Japan [22]. G9 deletion mutants were generated by PCR amplification or overlap extension PCR and inserted in frame with the MYC tag into the pCS2-MT-NLS expressing vector using the In Fusion Protocol (ST0345, Takara). All constructs were confirmed by Sanger sequencing.

### Cell Cultures and Co-Immunoprecipitation assays

Human embryonic kidney cells (HEK293T) were maintained in T-75 culture flasks at 37°C and 5% CO_2_. DMEM medium (Gibco) was supplemented with 10% fetal bovine serum (FBS, Gibco), 100 U/ml penicillin/streptomycin (Gibco) and 1mM sodium pyruvate (Gibco) (Maintenance Medium). Cells were subcultured when reaching 80 – 90% confluency. All media was replaced every 48 hours.

For transfection assays, HEK293T cells were plated on coated 10cm culture dishes (Greiner Bio-One, vented, sterile, PS coating) at a confluency of 3 x 10^6^ cells/dish. Dishes were used for plasmid transfection after reaching 50-80% confluency, usually 24h-48h after plating. 18-20 μg of indicated plasmids were transfected in HEK293T cells using the CalPhos Mammalian Transfection Kit (Takara). After 48h, cells were washed with RNAse Free ice-cold PBS and lysed for 15 – 20 mins in IPH Lysis Buffer (150 mM NaCl, 5 mM EDTA pH 8.0, 50 mM Tris pH8.0, 1% NP-40) containing protease inhibitor cocktail (cOmplete, EDTA-free Protease Inhibitor Cocktail – Roche). Supernatants were collected after centrifugation at 12000 RPM, 4°C and precleared by incubating with a mix of Protein G Plus/A Agarose Beads (Millipore) on a tube rotator for 2-3h, 4°C. Protein concentration was estimated with DC Protein Assay (Biorad) and equal amounts of protein (25 μg) were mixed with 5x Laemmli Buffer, heated up for 5 min, 95°C and run on 10% SDS-PAGE gels and transferred to nitrocellulose membranes (Amersham Protran Western blotting membranes – Sigma Aldrich) to validate protein quality and specificity.

48h after transfection, immunoprecipitations were performed with 1000 μg of total protein extracts from transfected cells and 5 μg of antibody overnight at 4°C under rotation. The day after, 40 μl of Protein A Sepharose CL-4B beads (Sigma; GE17-0780-01) was added in the tube and incubated for and additiona1 hour at 4°C under rotation. After three washes with IPH buffer 150 mM, immunoprecipitated proteins were eluted by heating at 100°C during 5 minutes in 1 x Laemmli Sample Buffer and subjected to Western blot analysis.

### Immunofluorescence

Dissected embryos were fixed in 4% Paraformaldehyde (PFA) for 15 minutes, washes 4 times with ice cold phosphate-buffered saline (PBS) and then, cryoprotected at 4^°^C overnight in 30% Sucrose dissolved in PBS. Embryos were embedded in 7,5% gelatin – 15% sucrose (dissolved in PBS) and stored at –80°C. The blocks are sectioned at the level of the thoracic dorsal root ganglia into 14μm sections at –30°C in the cryostat and collected slides are kept at –20°C. Immunostainings were performed as previously described [23] using mouse monoclonal α-H3K9me2 (Abcam, ab1220), mouse monoclonal α-G9a (R&D Systems, PP-A8620A-00), Goat polyclonal α-TrkA (R&D Systems, AF1056), Goat polyclonal α-TrkB (Cell Signaling, AF1494), Goat polyclonal α-TrkC (Cell Signaling, AF1404), Chicken polyclonal α-Peripherin (Abcam, ab106276), Rabbit polyclonal α-pH3 (Millipore, 07-690), Rabbit polyclonal cleaved α-Caspase (Cell Signaling, 9661), Rabbit polyclonal α-Phox2b (kind gift from Jean-François Brunet), Rabbit polyclonal α-Nav1.8 (Abcam, ab63331), Goat polyclonal α-Ret (R&D Systems, AF482), Rabbit polyclonal α-CGRP (Sigma-Aldrich, C8198), Mouse monoclonal α-Flag (Sigma-Aldrich, F1804), Rabbit polyclonal α-c-Myc (Sigma-Aldrich, PLA0001) and Normal Rabbit IgG (Cell Signaling, 2729). Secondary antibodies used in this study were Goat α-mouse IgG HRP (Jackson ImmunoResearch, 115-035-003), Goat α-rabbit IgG HRP (Cell Signaling, 7074), Goat α-mouse Alexa 594 (Invitrogen, A11032), Goat-α-rabbit Alexa Fluor 488 (Invitrogen, A11008), Goat α-rabbit Alexa Fluor 594 (Invitrogen, A11012), Goat α-guinea pig Alexa Fluor 488 (Invitrogen, A11073), Goat α-guinea pig Alexa Fluor 594 (Invitrogen, A11076), donkey α-mouse 488 (Invitrogen, A21202), donkey α-goat Alexa Fluor 594 (Invitrogen, A11058. Immunofluorescent images were acquired on a Zeiss Axio Observer Z1 fluorescent microscope or a laser-scanning confocal microscope Zeiss LSM 710 using the Zeiss Zen 2 microscope software.

### RT – qPCR

Total RNA from dissected E14.5 DRG was extracted using the Monarch Total RNA Miniprep Kit (New England Biolabs). cDNA was synthesized with iScript cDNA synthesis kit (Biorad) and RT-qPCR was performed using the Luna Universal qPCR Master Mix (New England Biolabs). The comparative 2^-ΔΔCT^ method was used to determine relative expression of the *G9a* cKO samples to compared the expression level of controls, overall normalized to *GAPDH* expression. The following primers were used: For *GAPDH*, forward primer 5’-CTCCCACTCTTCCACCTTCG –3’ and reverse 5’-GCCTCTCTTGCTCAGTGTCC –3’, for *G9a/Ehmt2* exons 24-25 forward primer 5’-GATCTACGGTTCCCACGCAT-3’ and reverse 5’-GTCACCGTAGTCAAAGCCCA-3’. Two-tailed Student’s t-test was used to measure statistical significance (* = p value <0.05, sample pool size n=4).

### Statistical Analysis

Statistical analyses were performed using GraphPad version 9. Cell countings were performed for at least 4 biological replicates per condition, on at least 8 DRG sections per sample. For statistical analysis, the unpaired, non-parametric Mann Whitney test was used (*= pvalue <0.05). Graphical quantifications were represented from qualitative data indicating the number (n) of embryos included in the analysis as individual values (mean +/-SEM).

---

**Figure S1.**
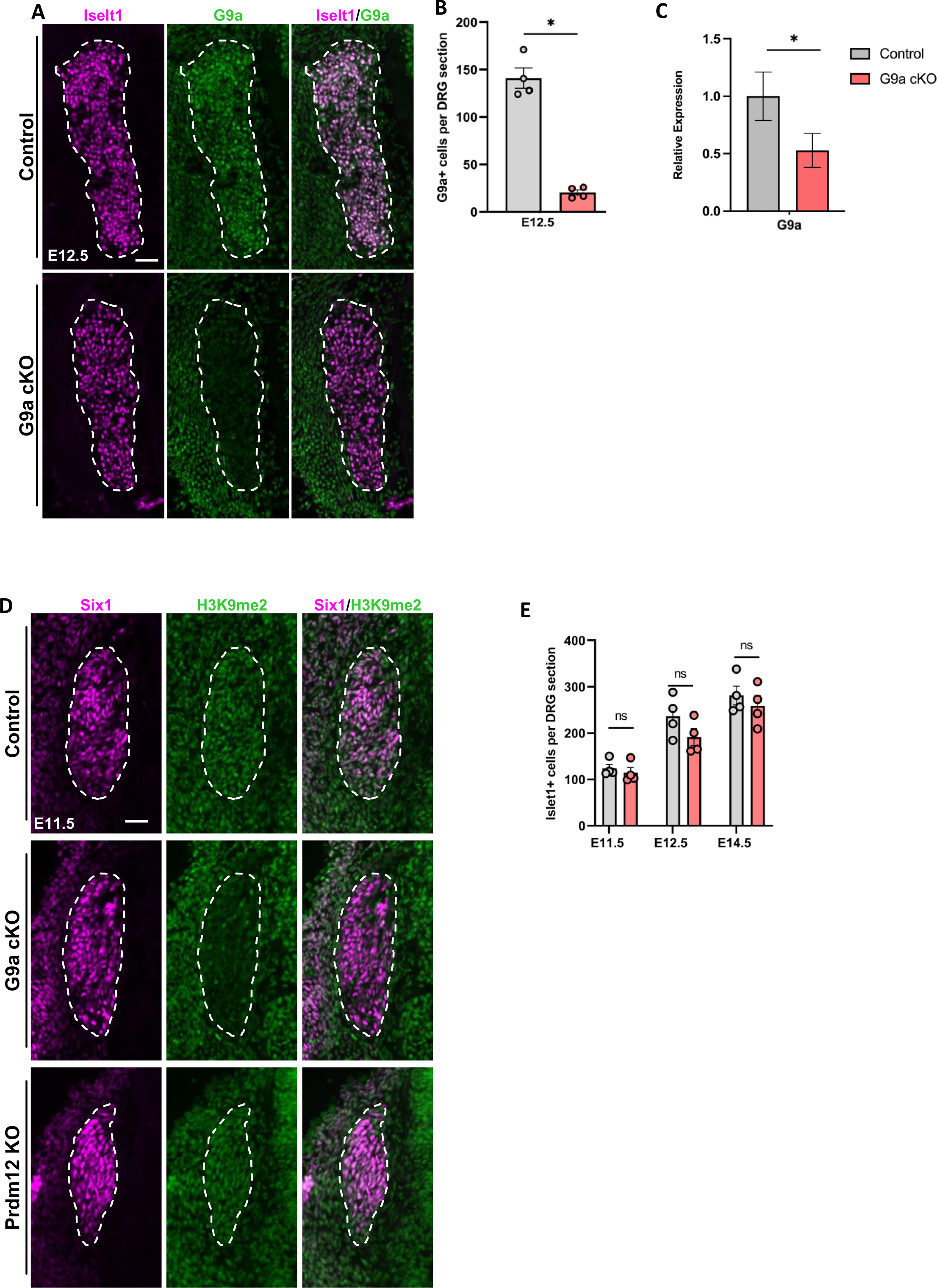
Validation of the G9a conditional knockout (cKO) mouse model. (A) Double immunostaining with the pan-sensory neuron marker Islet1 and G9a antibodies performed on coronal sections through dorsal root ganglia of control or *G9a* cKO embryos at E12.5. Scale bars, 50 μm. **(B)** Quantification of the mean number of G9a-positive cells on dorsal root ganglia coronal sections of control or *G9a* cKO embryos at E12.5. Each dot represents the mean value of G9a^+^ cells in one biological replicate. Mann-Whitney test. P-value, * < 0.005. Mean ± SEM **(C)** Relative expression of *G9a* quantified by RT-qPCR in DRG collected from E14.5 control and *G9a* cKO embryos. Mean ± SD, n=3. Student T test with Welch correction. P-value, * <0.005. **(D)** Quantification of the mean number of cells labelled with the pan-sensory neuron marker Islet1 in DRG coronal sections of control or *G9a* cKO embryos at indicated stages. Mann-Whitney test. P-value, ns > 0.999. **(E)** Double immunostaining with antibodies against Six1 and the histone methylation mark H3K9me2 on coronal sections through DRG of control, *G9a* cKO or *Prdm12* KO embryos at E11.5. Scale bars, 50 μm. DRG are delineated by white dashed lines.

**Figure S2.**
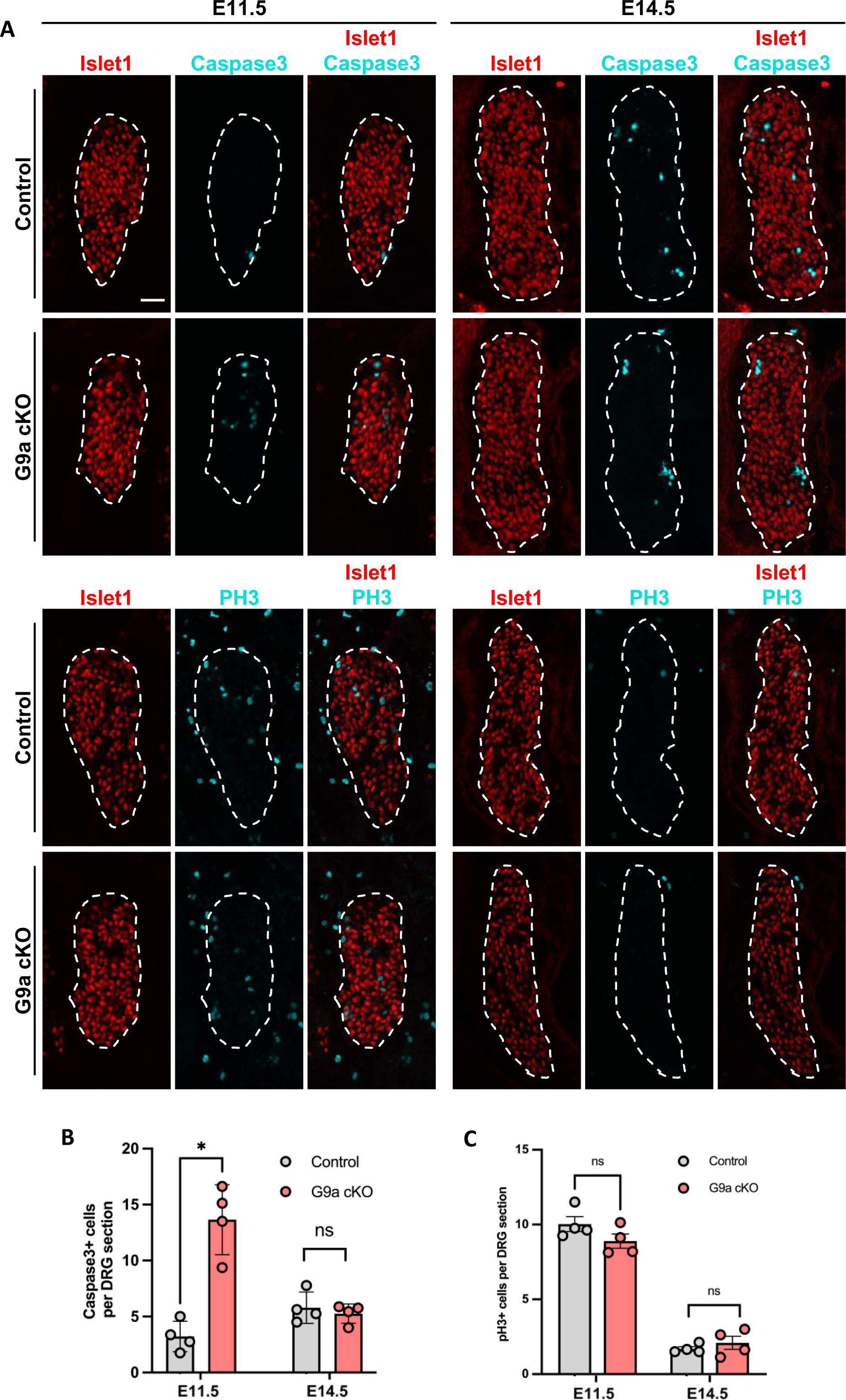
Loss of G9a results in a transient increased of apoptosis in DRG. (A) Double immunostainings with antibodies against the pan-neuronal marker Islet1 and the pro-apoptotic Cleaved-Caspase3 (upper panels) or against phospho-Histone H3 (PH3, lower panels) on coronal sections through DRG of control or *G9a* cKO embryos at E11.5 and E14.5. Scale bar, 50 μm. DRG are delineated by white dashed lines. **(B)** Quantification of the mean number of Caspase3^+^ cells on coronal sections through DRG of control or *G9a* cKO embryos at indicated embryonic stages. **(C)** Quantification of the mean number of PH3^+^ cells on coronal sections through DRG of control or *G9a* cKO embryos at indicated embryonic stages. Histograms are represented as mean ± SEM. Each dot represents the mean value obtained for an individual biological replicate. Mann-Whitney test. P-value, * <0.005, ns > 0.999.

**Figure S3.**
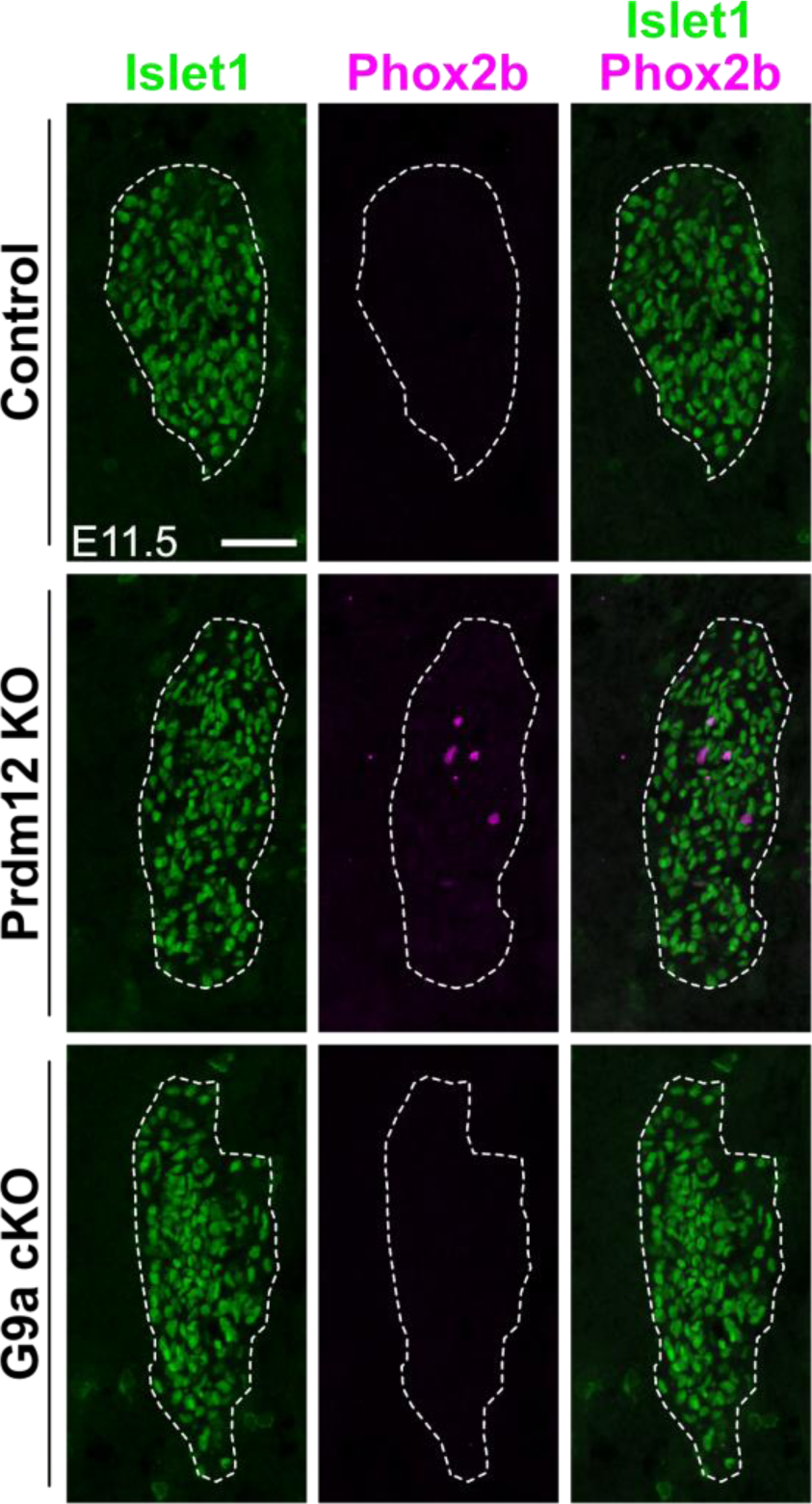
Loss of G9a is not accompanied by Phox2b ectopic expression. Double immunostainings with antibodies against the pan-neuronal marker Islet1 and the transcription factor Phox2b on coronal sections through DRG of control, *Prdm12* KO and *G9a* cKO embryos at E11.5. Scale bar, 50 μm. DRG are delineated by white dashed lines.

**Figure S4.**
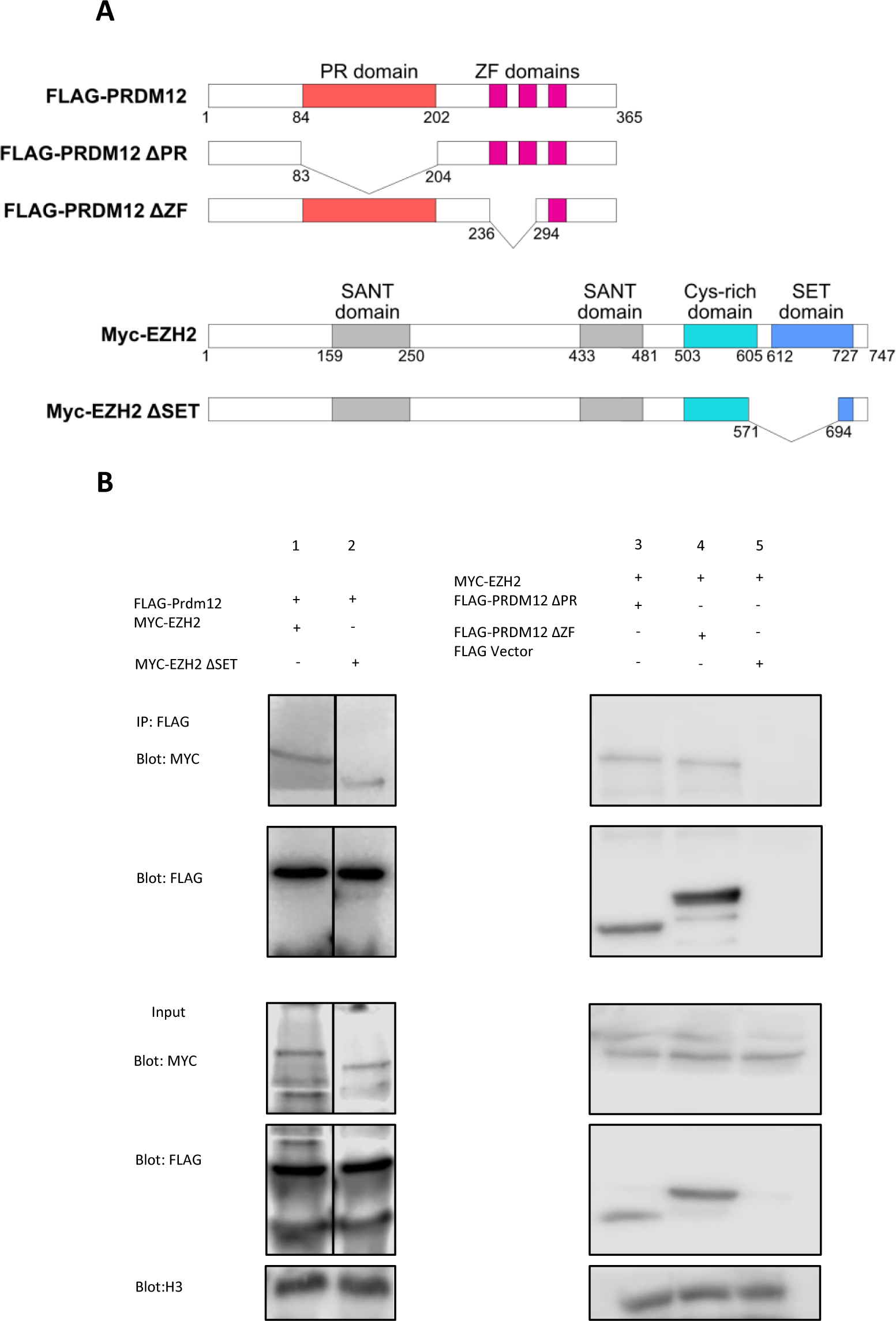
Prdm12 interacts with EZH2 independently of the zinc fingers and SET domains. (**A)** Schematic diagram of WT and deletion mutants of Flag-PRDM12 and Myc-EZH2, using an empty FLAG vector as a control. **(B)** HEK293T cells were transfected with the indicated plasmids. Lysates were immunoprecipitated with anti-Flag antibodies. Immunoprecipitates and 5% of the input were then subjected to western blot analysis with anti-Flag or anti-Myc antibodies.

## REFERENCES

1. Meltzer S, Santiago C, Sharma N, Ginty DD. The cellular and molecular basis of somatosensory neuron development. Neuron. 2021;

2. Marmigère F, Ernfors P. Specification and connectivity of neuronal subtypes in the sensory lineage. Nat Rev Neurosci. 2007;8:114–27.

3. Lallemend F, Ernfors P. Molecular interactions underlying the specification of sensory neurons. Trends Neurosci [Internet]. 2012;35:373–81. Available from: 10.1016/j.tins.2012.03.006

4. Vermeiren S, Bellefroid EJ, Desiderio S. Vertebrate Sensory Ganglia: Common and Divergent Features of the Transcriptional Programs Generating Their Functional Specialization. Front Cell Dev Biol. 2020;8.

5. Emery EC, Ernfors P. Dorsal Root Ganglion Neuron Types and Their Functional Specialization. The Oxford Handbook of the Neurobiology of Pain. 2020;127–55.

6. Vrontou S, Wong AM, Rau KK, Koerber HR, Anderson DJ. Genetic identification of C fibres that detect massage-like stroking of hairy skin in vivo. Nature. 2013;493:669–73.

7. Lou S, Pan X, Huang T, Duan B, Yang FC, Yang J, et al. Incoherent feed-forward regulatory loops control segregation of C-mechanoreceptors, nociceptors, and pruriceptors. Journal of Neuroscience [Internet]. 2015;35:5317–29. Available from: http://www.jneurosci.org/cgi/doi/10.1523/JNEUROSCI.0122-15.2015

8. Elias LJ, Succi IK, Schaffler MD, Foster W, Gradwell MA, Bohic M, et al. Touch neurons underlying dopaminergic pleasurable touch and sexual receptivity. Cell. 2023;

9. Sharma N, Flaherty K, Lezgiyeva K, Wagner DE, Klein AM, Ginty DD. The emergence of transcriptional identity in somatosensory neurons. Nature [Internet]. 2020;577:392–8. Available from: 10.1038/s41586-019-1900-1

10. Kokotović T, Langeslag M, Lenartowicz EM, Manion J, Fell CW, Alehabib E, et al. PRDM12 Is Transcriptionally Active and Required for Nociceptor Function Throughout Life. Front Mol Neurosci. 2021;14.

11. Bartesaghi L, Wang Y, Fontanet P, Wanderoy S, Berger F, Wu H, et al. PRDM12 Is Required for Initiation of the Nociceptive Neuron Lineage during Neurogenesis. Cell Rep. 2019;26:3484–3492.e4.

12. Landy MA, Goyal M, Casey KM, Liu C, Lai HC. Loss of Prdm12 during development, but not in mature nociceptors, causes defects in pain sensation. Cell Rep. 2021;34.

13. Desiderio S, Vermeiren S, Van Campenhout C, Kricha S, Malki E, Richts S, et al. Prdm12 Directs Nociceptive Sensory Neuron Development by Regulating the Expression of the NGF Receptor TrkA. Cell Rep [Internet]. 2019;26:3522–3536.e5. Available from: 10.1016/j.celrep.2019.02.097

14. Nagy V, Cole T, Van Campenhout C, Khoung TM, Leung C, Vermeiren S, et al. The evolutionarily conserved transcription factor PRDM12 controls sensory neuron development and pain perception. Cell Cycle [Internet]. 2015;14:1799–808. Available from: http://www.tandfonline.com/doi/full/10.1080/15384101.2015.1036209

15. Chen YC, Auer-Grumbach M, Matsukawa S, Zitzelsberger M, Themistocleous AC, Strom TM, et al. Transcriptional regulator PRDM12 is essential for human pain perception. Nat Genet [Internet]. 2015;47:803–8. Available from: 10.1038/ng0815-962b

16. Latragna A, Sabaté San José A, Tsimpos P, Vermeiren S, Gualdani R, Chakrabarti S, et al. Prdm12 modulates pain-related behavior by remodeling gene expression in mature nociceptors. Pain. 2021;

17. Imhof S, Kokotović T, Nagy V. PRDM12: New Opportunity in Pain Research. Trends Mol Med. 2020;26:895–7.

18. Hohenauer T, Moore AW. The Prdm family: Expanding roles in stem cells and development. Development (Cambridge). 2012;139:2267–82.

19. Fog CK, Galli GG, Lund AH. PRDM proteins: Important players in differentiation and disease. BioEssays. 2012;34:50–60.

20. Mzoughi S, Tan YX, Low D, Guccione E. The role of PRDMs in cancer: One family, two sides. Curr Opin Genet Dev [Internet]. 2016;36:83–91. Available from: 10.1016/j.gde.2016.03.009

21. Zannino DA, Sagerström CG. An emerging role for prdm family genes in dorsoventral patterning of the vertebrate nervous system. Neural Dev. 2015.

22. Yang CM, Shinkai Y. Prdm12 is induced by retinoic acid and exhibits anti-proliferative properties through the cell cycle modulation of P19 embryonic carcinoma cells. Cell Struct Funct [Internet]. 2013;38:195–204. Available from: http://www.ncbi.nlm.nih.gov/pubmed/23856557

23. Thélie A, Desiderio S, Hanotel J, Quigley I, Van Driessche B, Rodari A, et al. Prdm12 specifies V1 interneurons through cross-repressive interactions with Dbx1 and Nkx6 genes in Xenopus. Development (Cambridge) [Internet]. 2015;142:3416–28. Available from: http://dev.biologists.org/cgi/doi/10.1242/dev.121871

24. Danielian PS, Muccino D, Rowitch DH, Michael SK, McMahon AP. Modification of gene activity in mouse embryos in utero by a tamoxifen-inducible form of Cre recombinase. Curr Biol. 1998;8:1323– 6.

25. Tachibana M, Nozaki M, Takeda N, Shinkai Y. Functional dynamics of H3K9 methylation during meiotic prophase progression. EMBO J. 2007;26:3346–59.

26. Shankar SR, Bahirvani AG, Rao VK, Bharathy N, Ow JR, Taneja R. G9a, a multipotent regulator of gene expression. Epigenetics. 2013. p. 16–22.

27. Pattyn A, Morin X, Cremer H, Goridis C, Brunet J-F. The homeobox gene Phox2b is essential for the development of autonomic neural crest derivatives. Nature. 1999;399:366–70.

28. D’Autréaux F, Coppola E, Hirsch MR, Birchmeier C, Brunet JF. Homeoprotein Phox2b commands a somatic-to-visceral switch in cranial sensory pathways. Proc Natl Acad Sci U S A. 2011;108:20018–23.

29. Luo W, Wickramasinghe SR, Savitt JM, Griffin JW, Dawson TM, Ginty DD. A Hierarchical NGF Signaling Cascade Controls Ret-Dependent and Ret-Independent Events during Development of Nonpeptidergic DRG Neurons. Neuron. 2007;54:739–54.

30. Gyory I, Wu J, Fejér G, Seto E, Wright KL. PRDI-BF1 recruits the histone H3 methyltransferase G9a in transcriptional silencing. Nat Immunol. 2004;5:299–308.

31. Davis CA, Haberland M, Arnold MA, Sutherland LB, McDonald OG, Richardson JA, et al. PRISM/PRDM6, a transcriptional repressor that promotes the proliferative gene program in smooth muscle cells. Mol Cell Biol. 2006;26:2626–36.

32. Duan Z, Person RE, Lee H-H, Huang S, Donadieu J, Badolato R, et al. Epigenetic regulation of protein-coding and microRNA genes by the Gfi1-interacting tumor suppressor PRDM5. Mol Cell Biol. 2007;27:6889–902.

33. Biferali B, Bianconi V, Perez DF, Kronawitter SP, Marullo F, Maggio R, et al. Prdm16-mediated H3K9 methylation controls fibro-adipogenic progenitors identity during skeletal muscle repair. Sci Adv. 2021;7.

34. Higashihori N, Lehnertz B, Sampaio A, Underhill TM, Rossi F, Richman JM. Methyltransferase G9A Regulates Osteogenesis via Twist Gene Repression. J Dent Res. 2017;96:1136–44.

35. Ideno H, Nakashima K, Komatsu K, Araki R, Abe M, Arai Y, et al. G9a is involved in the regulation of cranial bone formation through activation of Runx2 function during development. Bone. 2020;137:115332.

36. Laumet G, Garriga J, Chen S-R, Zhang Y, Li D-P, Smith TM, et al. G9a is essential for epigenetic silencing of K(+) channel genes in acute-to-chronic pain transition. Nat Neurosci. 2015;18:1746–55.

37. Schwarz D, Varum S, Zemke M, Schöler A, Baggiolini A, Draganova K, et al. Ezh2 is required for neural crest-derived cartilage and bone formation. Development. 2014;141:867–77.

38. Rowitch DH, Echelard Y, Danielian PS, Gellner K, Brenner S, McMahon AP. Identification of an evolutionarily conserved 110 base-pair cis-acting regulatory sequence that governs Wnt-1 expression in the murine neural plate. Development. 1998;125:2735–46.

